## Supplementary Figures 1-9 for "An Interbacterial Cysteine Protease Toxin Inhibits Cell Growth by Targeting Type II DNA Topoisomerases GyrB and ParE"

### Supplementary Figure 1

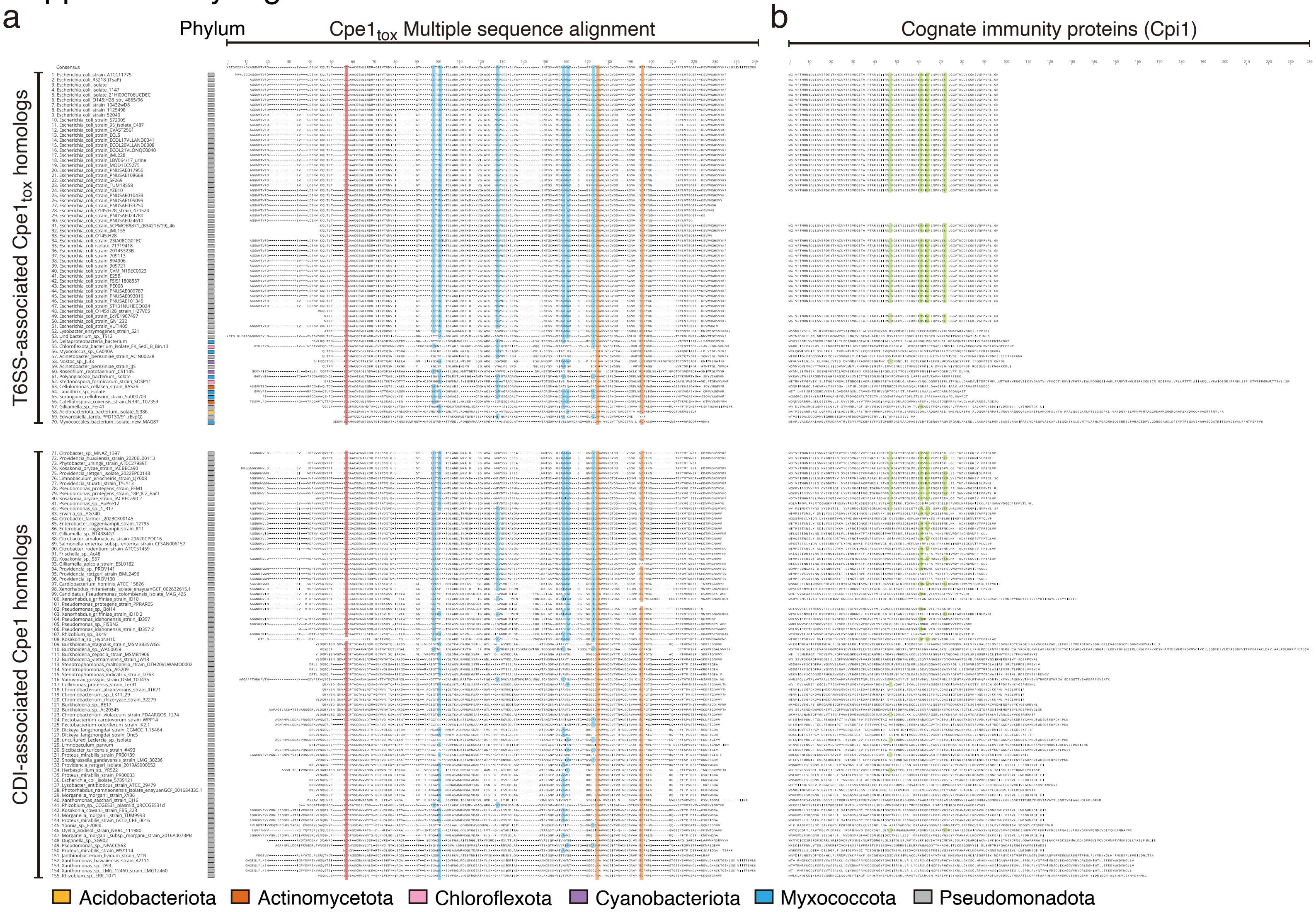

**Supplementary Figure 1. Interbacterial PLCP effectors and their cognate immunity proteins are distributed across various Gram-negative bacteria.** Related to **Figure 1b. (a)** Multiple sequence alignment of 155 unique sequences of interbacterial PLCP effectors. Sequences have been ordered by relative identity to $Cpe1_{tox}$ of *E. coli* ATCC-11775 and are displayed in two clusters divided according to the associated secretion systems (T6SS or CDI). Conserved residues are indicated as follows: the catalytic triad residues are highlighted by red (Cysteine) and orange (Histidine and Aspartate/Asparagine) shadows; and hydrophobic residues composing the surface concave region are highlighted by blue shadows. The Phylum of each strain containing the interbacterial PLCP effectors is indicated by colored boxes (as indicated in the legend). **(b)** Sequences of the putative cognate immunity proteins related to the interbacterial PLCP effectors in **(a)**. Conserved binding residues are highlighted in green shadows. Blank rows indicate an absence of putative immunity proteins due to incomplete genomic data for certain bacterial species in the NCBI database. See also **Supplementary Table 1** for detailed information.

### Supplementary Figure 2

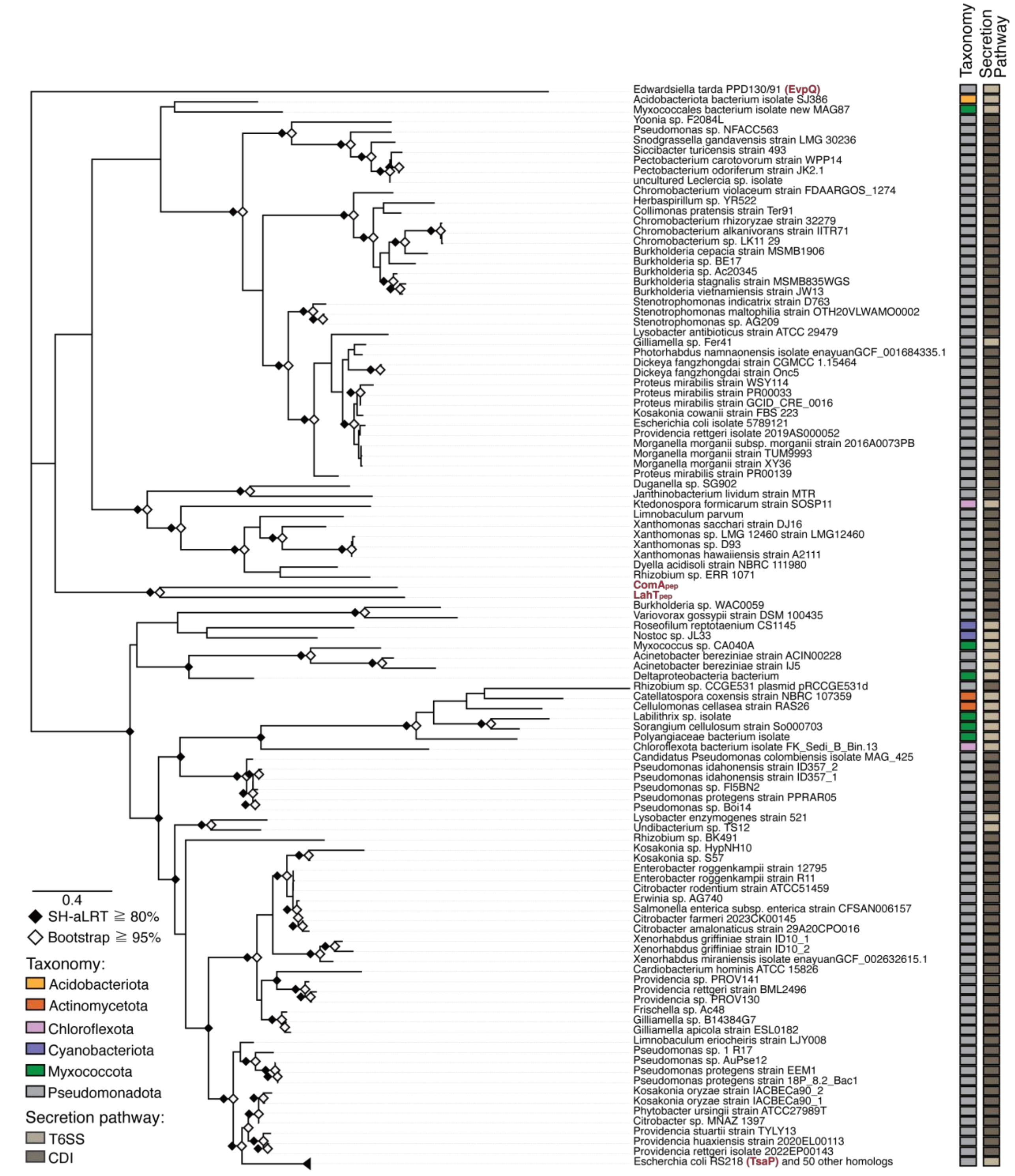


**Supplementary Figure 2. Interbacterial PLCP effectors exhibit no correlations in phylogeny, associated secretion pathways, or taxonomy.** Related to **Figure 1d**. Maximum likelihood phylogeny of 155 unique PLCP effector sequences visualized as a rectangle tree. Branch support values (SH-aLRT and ultrafast bootstrap) are displayed as symbols (filled diamond: SH-aLRT ≥ 80%, hollow diamonds: ultrafast bootstrap ≥ 95%). Tips representing sequences of the closely-related housekeeping PLCPs, ComA$_{pep}$ and LahT$_{pep}$, as well as the previously identified interbacterial PLCP effectors EvpQ and TsaP, are highlighted in red. The taxonomy of strains containing the PLCP effectors and their associated secretion pathways is indicated on the right as colored boxes. The scale bar represents the average number of substitutions per site. See also **Supplementary Table 1** for detailed information.

Supplementary Figure 3

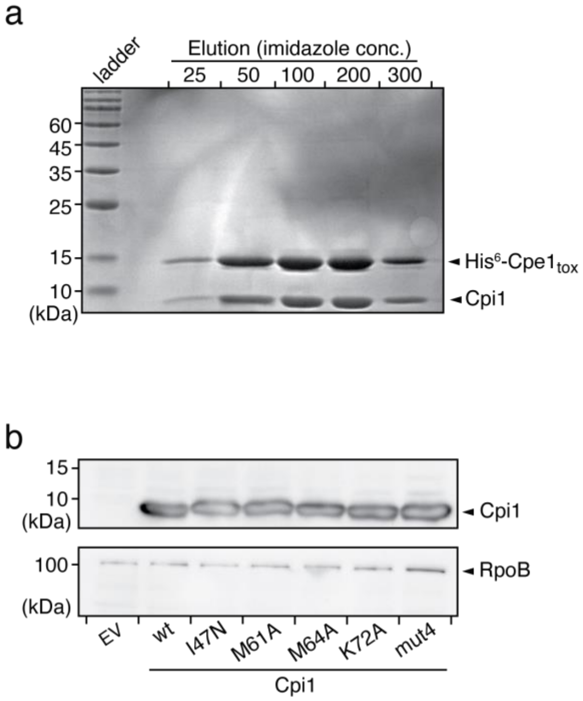


**Supplementary Figure 3. Purification of the Cpe1$_{tox}$-Cpi1 complex and protein expression levels of Cpi1 variants.** Related to **Figure 2. (a)** Coomassie-stained SDS-PAGE analysis of the His-tagged Cpe1$_{tox}$ co-purified with Cpi1. **(b)** Protein expression levels of wild-type Cpi1 and variants, as assessed by immunoblotting analysis. The cytosolic protein RpoB was used as a loading control.

Supplementary Figure 4

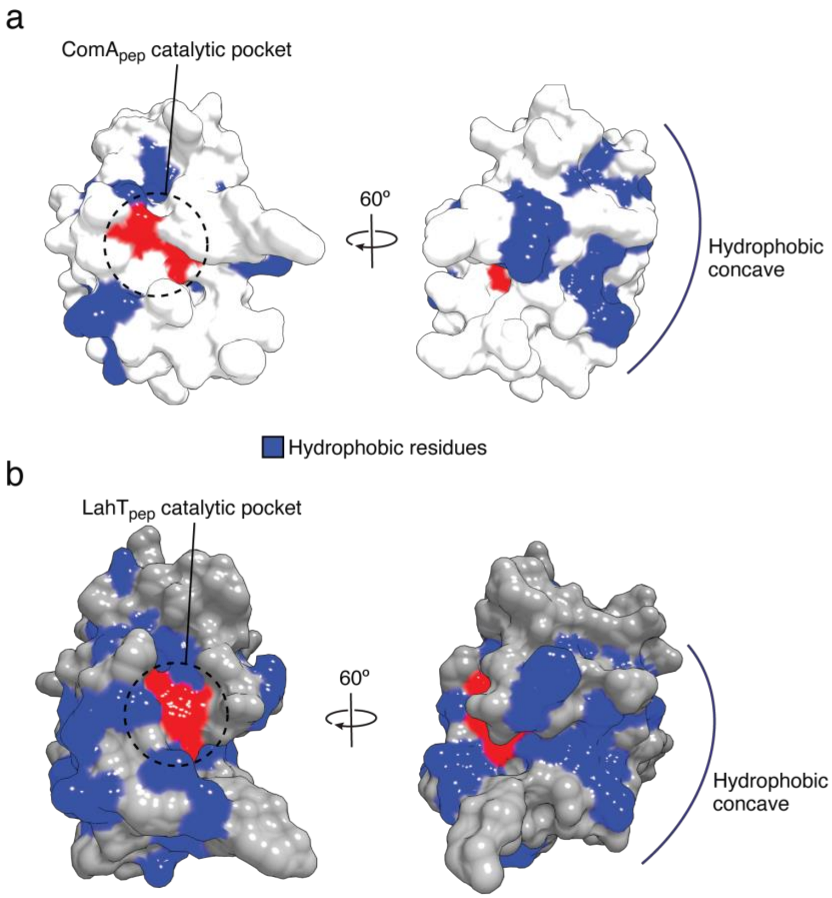


**Supplementary Figure 4. ComA$_{pep}$ and LahT$_{pep}$ feature hydrophobic concave regions on their surfaces near the catalytic pocket.** Related to **Figure 3. (a, b)** Depiction of ComA$_{pep}$ **(a)** and LahT$_{pep}$ **(b)** illustrating the location of their respective catalytic pockets and hydrophobic concave surfaces. ComA$_{pep}$ (PDB: 3K8U, light gray). LahT$_{pep}$ (PDB: 6MPZ, dark gray).

#### Supplementary Figure 5

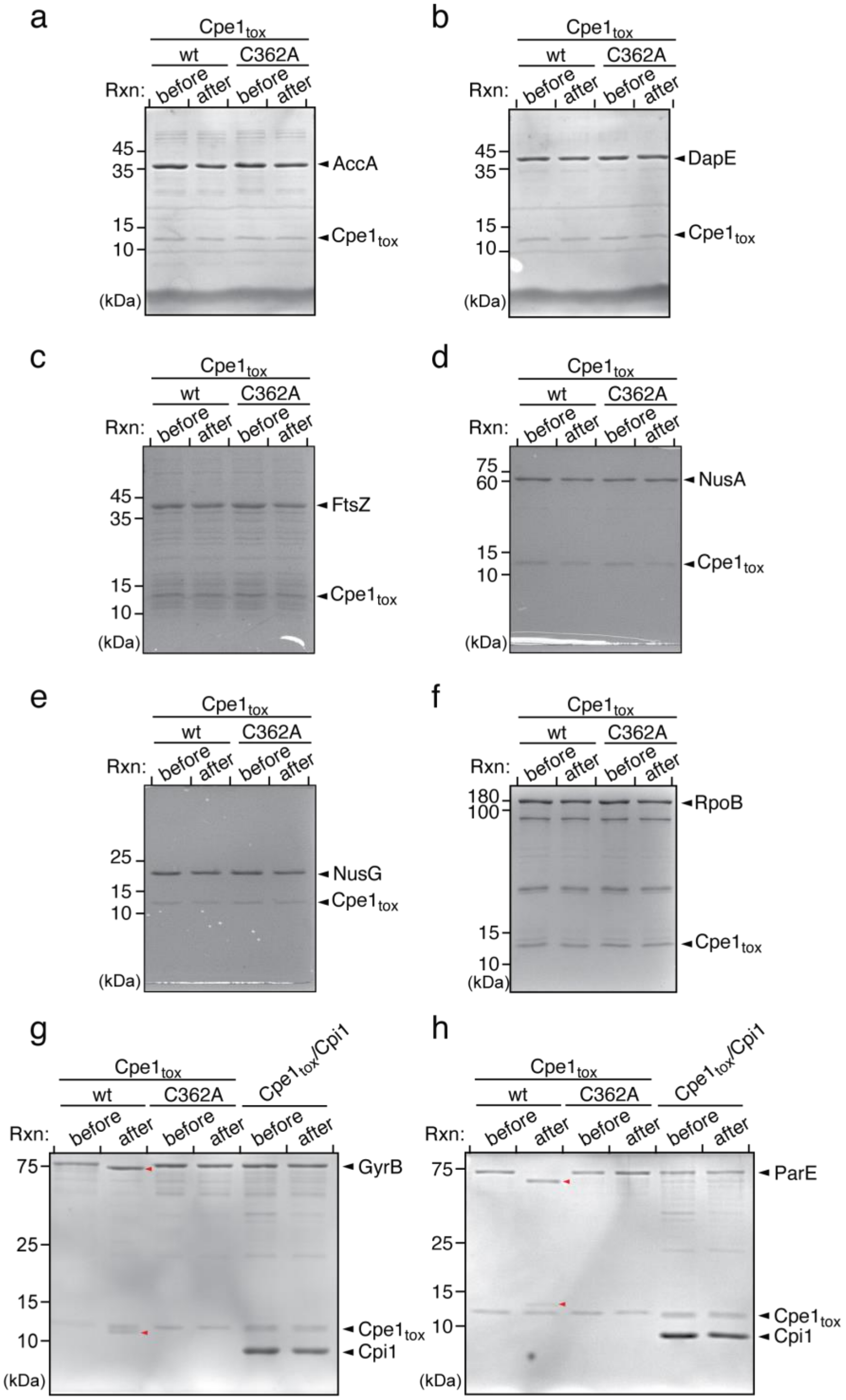


**Supplementary Figure 5. Cpe1$_{tox}$ specifically targets and cleaves GyrB and ParE among eight candidate substrates identified from the interactome analysis.** Related to **Figure 3. (a-f)** Coomassie-stained SDS-PAGE analysis of the *in vitro* cleavage assay of six candidate substrates of Cpe1$_{tox}$: AccA **(a)**, DapE **(b)**, FtsZ **(c)**, NusA **(d)**, NusG **(e)**, and RpoB **(f)**. **(g, h)** Presence of Cpi1 suppressed the cleavage of GyrB **(g)** and ParE **(h)** by Cpe1$_{tox}$. The substrates were incubated with Cpe1$_{tox}$ (lanes 1 and 2), Cpe1$_{tox}^{C362A}$ (lanes 3 and 4), or Cpe1$_{tox}$ and Cpi1 (lanes 5 and 6). Cleaved fragments are indicated with red arrowheads.

Supplementary Figure 6

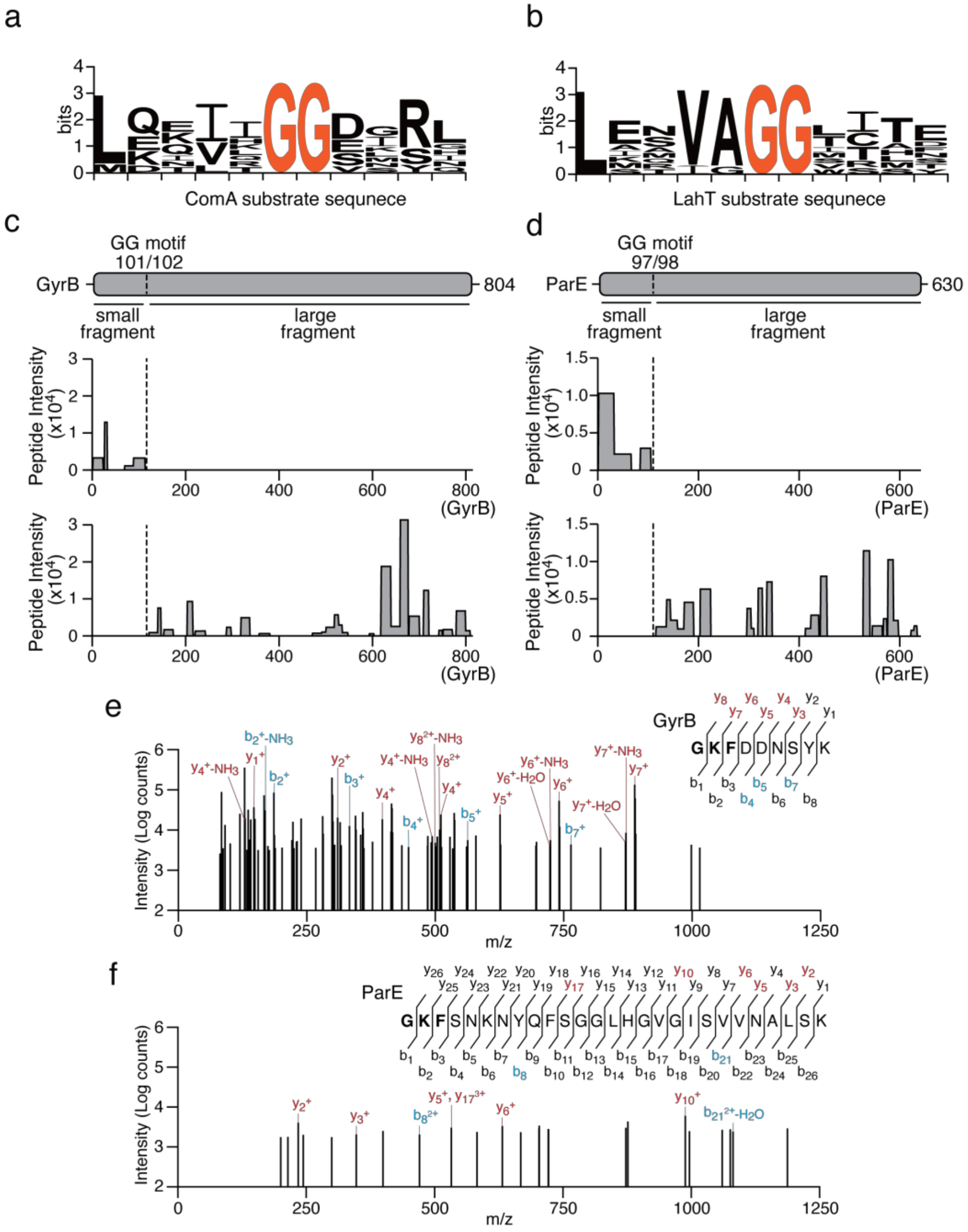


**Supplementary Figure 6. Cleavage sequences of ComA and LahT and mass spectrometric analysis of Cpe1-cleaved fragments.** Related to **Figure 4. (a, b)** Sequence logos showing the consensus sequence recognized by ComA **(a)** and LahT **(b)**. **(c, d)** Mapping of peptide sequences from Cpe1$_{tox}$-digested GyrB **(c)** and ParE **(d)** fragments. Cpe1$_{tox}$-cleaved fragments, resolved by Coomassie-stained SDS-PAGE, were purified, trypsin-digested, and subjected to MALDI-TOF analysis. The intensity of the signals (Y-axis) from identified peptides and their coverage across the respective full-length protein (X-axis) are plotted below. Upper chart: peptides from the small fragment. Lower chart: peptides from the large fragment. **(e, f)** Tandem mass spectrum of indicated peptides from Cpe1tox-cleaved GyrB **(e)** and ParE **(g)** fragments. Fragmentation ions (b, blue; y, red) with resolved spectra and the residues correlating to the LHAGGKF motif (bold) are indicated.

#### Supplementary Figure 7

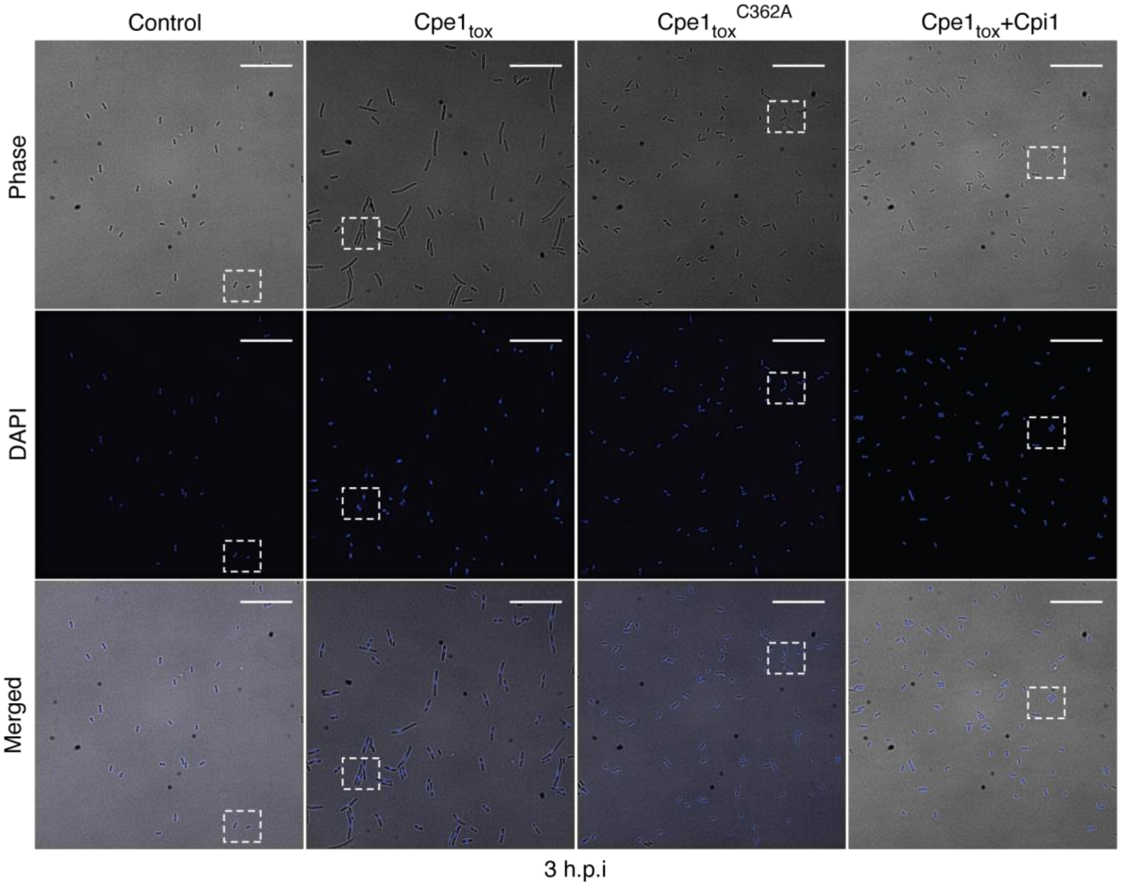


**Supplementary Figure 7. Cpe1 intoxication inhibits chromosome segregation in *E. coli*.** Related to **Figure 4f. (a)** Phase-contrast (top), blue fluorescence (middle), and merged (bottom) images of *E. coli* carrying an empty vector, *E. coli* expressing $Cpe1_{tox}$, *E. coli* expressing $Cpe1_{tox}^{C362A}$, or *E. coli* co-expressing $Cpe1_{tox}$ and Cpi1, after three hours of induction. Scale bar = 20 µm. The white borders demarcate the cropped images displayed in **Figure 4f**.

#### Supplementary Figure 8

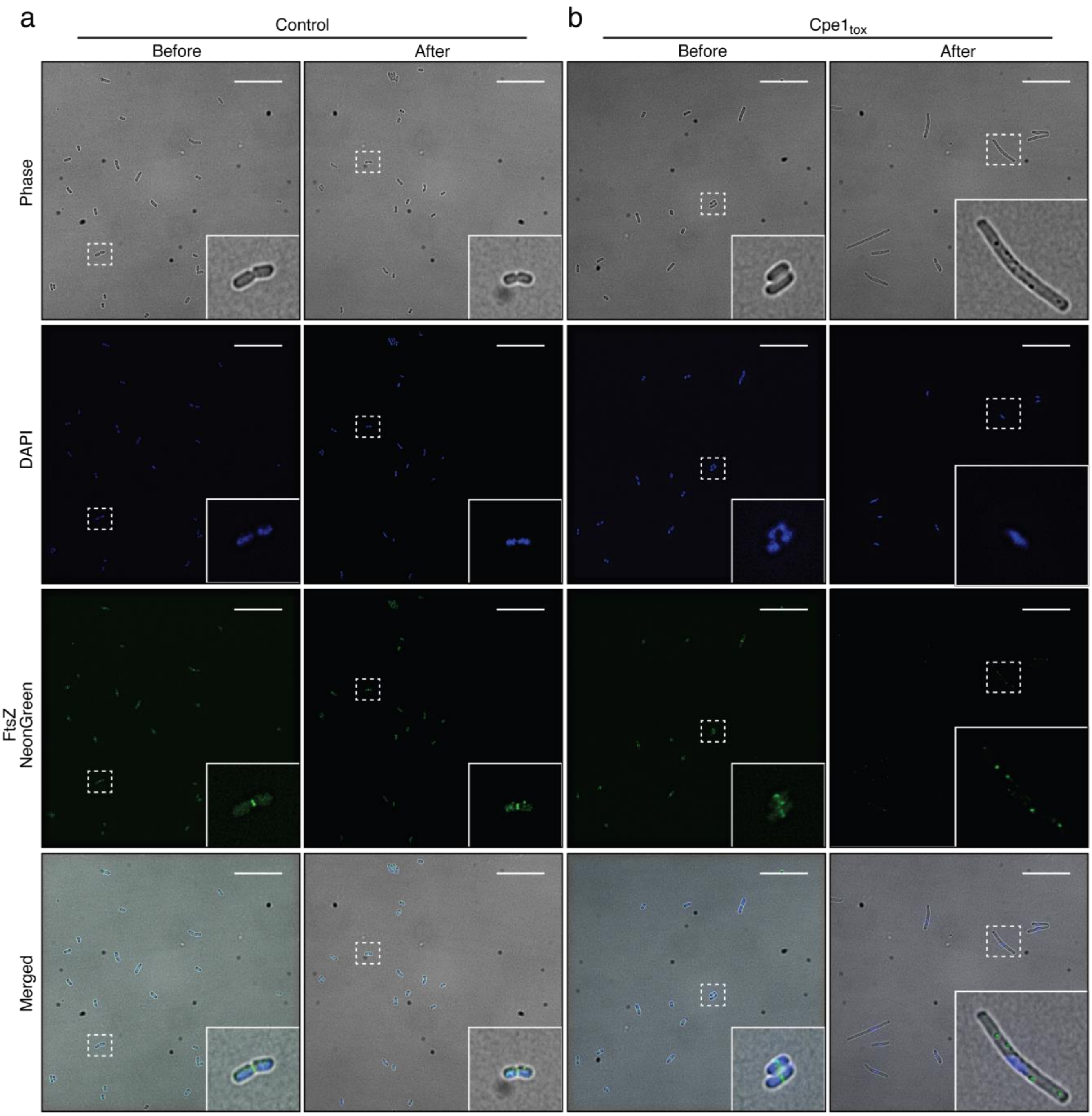


**Supplementary Figure 8. Z-ring formation is disrupted in Cpe1-intoxicated *E. coli*.** Related to **Figure 4f.** **(a)** Phase-contrast (top), blue fluorescence (second from top), green fluorescence (third from top), and merged (bottom) images of *E. coli* carrying an empty vector, shown before and after 2-hour incubation. **(b)** Fluorescence micrographs of *E. coli* intoxicated by $Cpe1_{tox}$. Phase-contrast (top), blue fluorescence (second from top), green fluorescence (third from top), and merged (bottom) images are presented. White borders indicate the zoomed-in regions shown in the bottom-right corner of each image. Scale bar = 20 μm.

#### Supplementary Figure 9

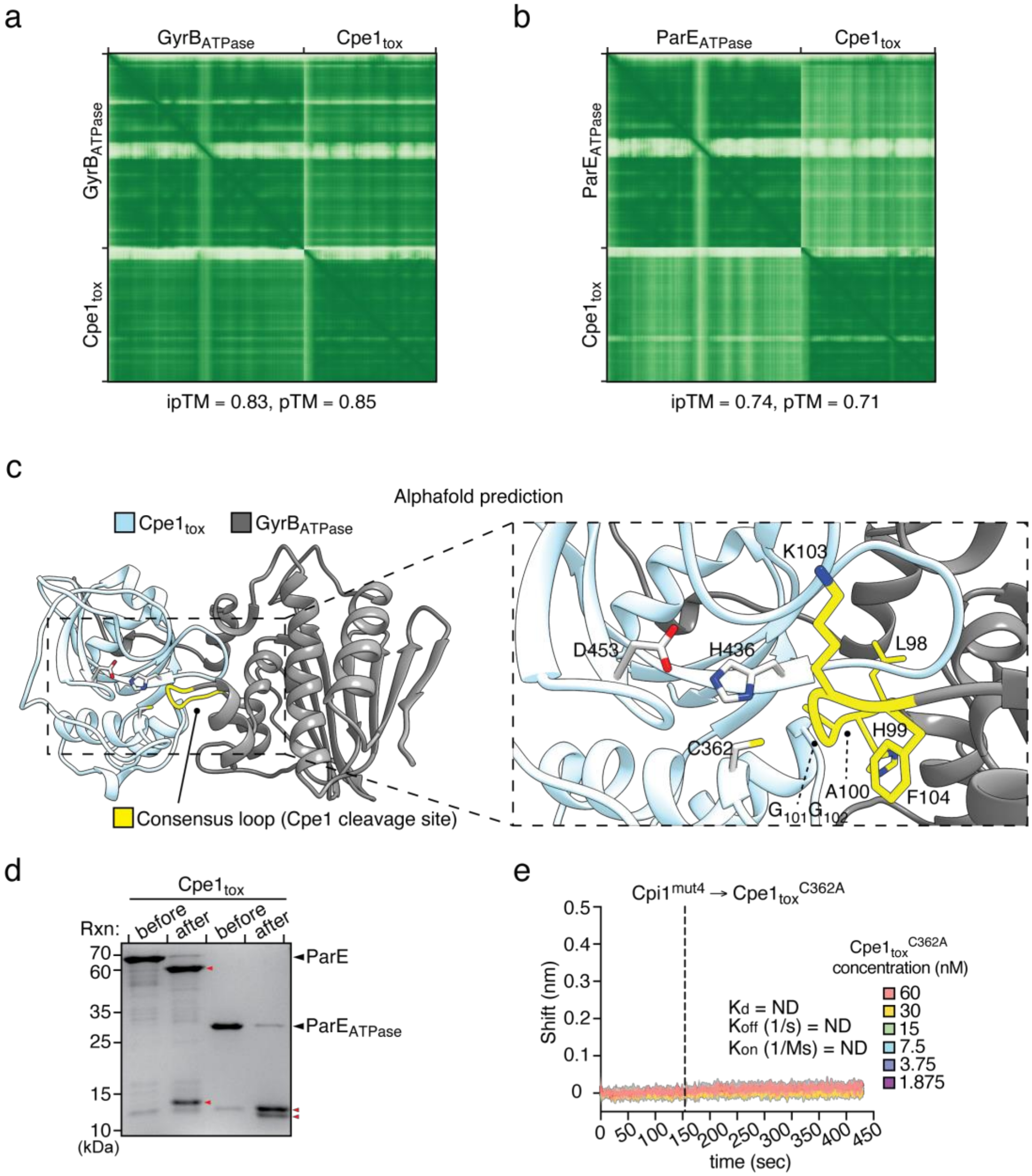


**Supplementary Figure 9. Interactions between $Cpe1_{tox}$ with the substrates and kinetics of $Cpe1_{tox}$ with $Cpi1^{mut4}$.** Related to **Figures 5 a, e, f. (a, b)** Predicted assigned error (PAE) plots for the models of the $Cpe1_{tox}$-$GyrB_{ATPase}$ interaction **(a)** and the $Cpe1_{tox}$-$ParE_{ATPase}$ interaction **(b)** in **Figure 5a**. Accuracy of the predicted relative positions of subunits within the complex, as indicated by AlphaFold 3 prediction scores (iPTM). Confidence in the overall folding of the complex is indicated by pTM scores. **(c)** Predicted structure of $Cpe1_{tox}$-$GyrB_{ATPase}$ interaction, showing the entrance of the consensus loop of GyrB into the active site of Cpe1 (left panel). The magnified view of the active pocket (right panel) implies possible catalysis. **(d)** $Cpe1_{tox}$ targets the ATPase domain of ParE. Cleavage of full-length ParE and the ATPase domain by $Cpe1_{tox}$ was analyzed by Coomassie-stained SDS-PAGE. Cleaved fragments are indicated with red arrowheads. **(e)** Results of kinetic assays on interactions between $Cpi1^{mut4}$ and $Cpe1_{tox}^{C362A}$ using biolayer interferometry (BLI). Data from three replicates have been plotted on an X-Y scatter graph. The Y-axis represents wavelength shift (nm) generated by binding of the two proteins, which is nearly non-detectable (N.D.). The X-axis represents reaction time in seconds. A vertical dashed line marks the transition from the association step to the dissociation step. Kinetic parameters such as $K_d$, $K_{on}$, $K_{off}$, and Rmax could not be calculated and are labeled as ND.
